# Improving MHC Class I antigen processing predictions using representation learning and cleavage site-specific kernels

**DOI:** 10.1101/2021.10.17.464741

**Authors:** Patrick J. Lawrence, Xia Ning

**Affiliations:** Biomedical Informatics Department, The Ohio State University, Columbus, OH, US; Computer Science and Engineering Department, The Ohio State University, Columbus, OH, US; Translational Data Analytics Institute, The Ohio State University, Columbus, OH, US

**Keywords:** artificial intelligence, machine learning, immunology, antigen processing, MHC Class I

## Abstract

In this work, we propose a new deep learning model, `MHCrank`, to predict the probability that a peptide will be processed for presentation within the MHC Class I pathway. We find that the performance of our model is significantly higher than two previously published baseline methods: `MHCflurry` and `netMHCpan`. Gains in performance result from the utilization of cleavage site-specific kernels and learned representations for amino acids. By visualizing the site-specific amino acid enrichment among top-ranked peptides, we find MHCrank’s top-ranked peptides are enriched at biologically relevant positions with amino acids that are consistent with previous work. Furthermore, the cosine similarity matrix derived from MHCrank’s learned embeddings for amino acids correlate highly with physiochemical properties that have been experimentally shown to be important in determining a peptide’s favorability to be processed. Altogether, the results reported in this work indicate that the proposed MHCrank demonstrates strong performance compared to existing methods and could have vast applicability to aid drug and vaccine development.

## Introduction

The major histocompatibility complex (MHC) Class I protein is a vital part of the immune system’s response to intracellular invasion by viruses, bacteria and parasites and against tumorigenesis.^1^ Its primary responsibility is to present antigens – short peptides eight to ten amino acids in length that are cleaved from proteins – into the extracellular environment to be recognized by cytotoxic (CD8^+^) T cells, which subsequently eliminate compromised cells via apoptosis.^2^ Thus, these peptides can be leveraged for the development of both vaccines that prime CD8^+^ T cells against a pathogen and drugs that elicit cytolytic activity in tumor cells.

There is not a single MHC Class I molecule. Rather, multiple versions can be produced based on the human leukocyte antigen (HLA) alleles present in an individual’s genome. HLA is the portion of the MHC class I molecule that binds presented peptides; hypermutability in HLAs’ binding groove yields variability in the binding affinity of processed peptides and affords greater coverage of the number of pathogens that can be recognized.^3^

The peptides which are presented by MHC Class I molecules must first undergo a series of processing steps to make the peptide more favorable for presentation. Peptidases digest proteins into fragments based on identified consensus sequences that indicate where to cleave proteins.^4^ Fragments from digested protein are then translocated across the rough endoplasmic reticular membrane by the TAP protein.^5, 6^ TAP filters these peptides based on which are most likely to have high affinity for the MHC Class I molecule. Specifically, TAP has a higher affinity for peptides between 8–16 amino acids in length,^7^ as well as peptides with either hydrophobic or basic C-terminal amino acids.^2, 5^ Once in the RER, longer peptides may be further cleaved from the N-termini to optimize its binding affinity,^4^ but the C-terminus remains untouched as this is the primary anchor point between the antigen and MHC molecule. Thus, leveraging these peptides for vaccine and drug development requires an understanding on which peptides will have the greatest opportunity to bind to any given HLA allele.

As a result, computer-aided methods have been developed to identify candidate peptides.^1, 8–12^ Among them, deep learning (DL) models that rank peptides’ binding affinities to MHC class I molecule(s) have achieved superior performance.^9^ These models are created with the goal of predicting which peptides will have the highest binding affinity for the HLA alleles. However, there is no guarantee that highly ranked peptides will be selected for presentation by upstream proteins,^8^ meaning that these models may lack biological relevance. Recent attempts have been made to develop models that rank the likelihood of peptides being processed within the MHC class I presentation pathway.^8, 9^ These are considered HLA-independent as they do not require any information about the HLA alleles, making them a more generalized approach. By training on peptides that have been confirmed to be processed for and presented by MHC Class I molecules, in combination with the amino acid residues immediately flanking the peptides in their original protein, such models can–in theory–learn the features that make a peptide more likely to be cleaved and processed for presentation. This incorporates the biological information missing from binding affinity models and has the potential to enable superior performance on predicting presentation.

In this work, we propose a novel DL, antigen processing (AP) prediction model, denoted `MHCrank`, that has been developed to rank candidate peptides by their likelihood to be processed for MHC Class I presentation. Based on the architecture used by O’Donnell *et al*,^8^ our model imparts additional biological relevance, focusing on the carboxyl (C)-terminal cleavage site of the antigen and pre-processing antigen sequences to simulate what is observed *in vivo*.^8^ In our development of `MHCrank`, we elect to forgo the use of the widely used BLOSUM62 matrix for amino acid representations. Instead, `MHCrank` learns a problem-specific embedding for each amino acid. Our experiments on the benchmark data set demonstrate `MHCrank` achieves a significant performance improvement over the compared baseline methods: `netMHCpan-4.0 eluted ligand` and `MHCflurry-2.0 antigen processing`, denoted `netMHCpan-EL` and `MHCflurry-AP`, respectively.

## Materials

### Data

We used the exact data sets as those used in O’Donnell *et al*.^8^ for our training, validation and testing of `MHCrank`. Specifically, we used the data which was employed to create and evaluate their `MHCflurry-AP` predictor. A more comprehensive description of the data is present below:

#### Training data

Our training data set was identical to that which was used by O’Donnell *et al*. to train their `MHCflurry-AP` predictor.^8^ This data set was compiled from two studies^13, 14^ and comprised the aggregate data from 100 mass spectroscopic (MS) experiments, including measurements on 8,537,960 distinct peptides and 92 different HLA alleles. Based on the nature of the MS experiments, bound peptides (hits) must have first underwent processing for MHC Class I presentation. Hits with a sequence length between eight and fifteen amino acids were selected as this range is optimal for presented peptides. O’Donnell *et al*. randomly generated 99 negative decoys per hit. Each decoy was the same length and was extracted from the same protein as the hit to which it corresponds. O’Donnell *et al*. used their binding affinity predictor to select decoys most similar to hits in terms of their predicted binding affinities. The hits and decoys selected were within the top 2% of predicted binding affinities for hits and decoys, respectively. The exclusion of weak binding hits and decoys from the data set was aimed at facilitating the model’s ability to learn features that strongly influence a peptide’s likelihood to be processed for presentation as opposed to learn features associated with binding affinity. This yielded 399,392 peptides in the training data, 44% of which were hits.^8^ The data was split into 4 training subsets (folds) by randomly withholding 10 MS experiments from each. As a result, the number of samples in each fold was 365,746; 352,144; 361,864 and 358,374, respectively. Additionally, each fold had 10% of its samples randomly withheld for validation.

#### Testing data processing

We used the same testing set to evaluate our `MHCrank` model as was used by O’Donnell *et al*. to evaluate their model.^8^ The data in the testing data set was compiled from 2 studies published in 2019; these studies comprised 20 experiments and identified 27,007 binding peptides.^14, 15^ According to O’Donnell *et al*., these specific experiments were withheld for testing as they were not yet published when their baseline methods, such as `netMHCpan`, were created.^8^ Therefore, none of the models being evaluated would have been exposed to the testing data for their training. 99 decoys were introduced for each hit in the testing set, bringing the total number of testing peptides up to 2,700,700. The testing set was also used by O’Donnell *et al*. to benchmark their binding affinity predictors. This required each peptide to be paired with multiple, distinct HLA alleles. However, for HLA-independent models, such as `MHCrank` and our baselines, which do not use HLA allele information, these distinct combinations are instead interpreted as duplicated samples that achieve identical scores. As a result, duplicated samples dominate the results and can either negatively or positively bias ranking performance. To prevent this, we removed any duplications of a peptides from the test set, leaving a remaining 2,409,183 peptides. Approximately, 0.73% of the remaining peptides were hits.

### Baseline methods

`MHCrank` was compared to two baseline methods: `MHCflurry-AP` and `netMHCpan-EL`. Both the baselines, similar to `MHCrank`, are AP prediction models. `MHCflurry-AP` was selected for its high performance. Note that because `MHCrank` architectures were leveraged from `MHCflurry-AP`, by comparing `MHCrank` to `MHCflurry-AP`, any improvements in performance exhibited by `MHCrank` may be associated with the introduced architectural alternations. The other baseline, `netMHCpan-EL`, was selected as it was identified by O’Donnell *et al*. to be the best performing AP predictor available.^8^ The predictions we used to evaluate the performance of `MHCflurry-AP` and `netMHCpan-EL` were those reported by O’Donnell *et al*.^8^

## Methods

The `MHCrank` architecture is presented in Figure 1b. The proposed `MHCrank` takes three types of information as input: 1) uniform-length sequence N-flank+peptide+C-flank, 2) C-flank cleavage site sequence, and 3) the peptide’s original length. The N- and C-flanks are defined as the first five amino acids adjacent to a peptide on its amine (N-) and carboxyl (C-) terminal ends, respectively.

**Figure 1:**
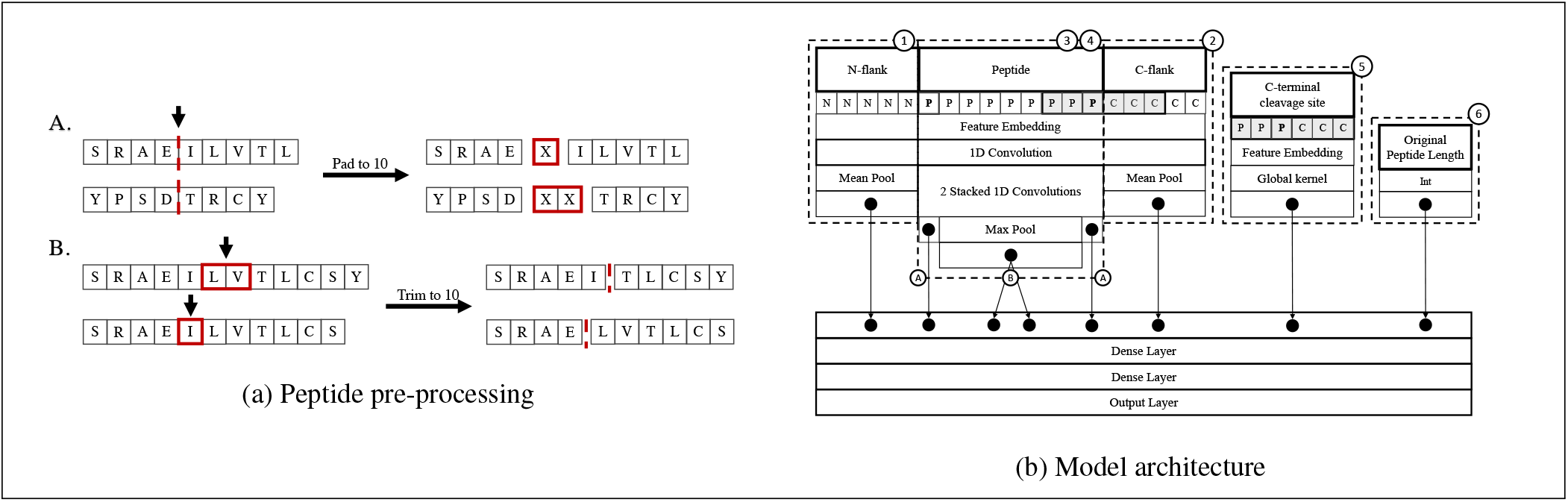
Peptide processing and model architecture. **Figure 1a**: Peptides undergo pre-processing to ensure they all possess a uniform length. (**A**) Peptides shorter than the desired length are padded with a number of ambiguous amino acid ’X’ to their center. Raw peptides with an odd length have padding offset from their center by one amino acid towards the N-flank. (**B**) Peptides longer than the desired length are trimmed at their center. Trimming peptides from an odd-length to an even-length and vice versa requires the trim to be offset from their center by one amino acid towards the N-flank. **Figure 1b**: `MHCrank` takes a uniform-length N-flank+peptide+C-flank sequence, C-terminal cleavage site (see gray box), and the peptide’s original length before padding or trimming as input. The amino acids comprising the sequence and `CSSK` undergo feature embedding. A convolution layer is applied to the embedding of the entire sequence. The remainder of the `MHCrank` architecture can be split into six components. Component (**1**) applies a mean pool to the convolution output corresponding to the N-flank. Component (**2**) applies a mean pool to the convolution output corresponding to the C-flank. The convolution output corresponding to the peptide sequence is forwarded to 2 stacked convolution layers. Components (**3**) and (**4**) each have 2 outputs (**A** and **B**) obtained from the output of these convolution layers. (**3A**) extracts the output corresponding to the peptide’s N-terminal amino acid. (**4A**) extracts the output corresponding to the peptide’s C-terminal amino acid. (**3B**) applies a mean pool to the peptide’s non-N-terminal amino acids. (**4B**) applies a mean pool to the peptide’s non-C-terminal amino acids. Component (**5**) applies a global kernel to the embedded `CSSK`. Component (**6**) is a single node that takes the peptide’s original length as input. Two dense layers are applied to the concatenated output of each component. The output from the second dense layer enters an output layer that predicts the probability of the input peptide undergoing antigen processing.

The C-flank cleavage site sequence is comprised of the terminal *r* amino acids of the peptide and the initial *r* amino acids of the C-flank. In the following subsections, we present how `MHCrank` learns from a peptide and its N- and C-flanks to predict antigen processing.

### Peptide pre-processing

As in other methods, all input N-flank+peptide+C-flank sequences are first processed to obtain uniformity in length before being passed to `MHCrank`. Peptides of various lengths (8-15 amino acids as in our data set) can bind to MHC Class I molecules because only their termini interact with and anchor to the molecules^2^. The central amino acids of a bound peptide create an arch-like formation as the peptide bows increasingly away from the pocket with increased length.^16^ As the central amino acids do not contribute directly to binding, their inclusion in `MHCrank` may be uninformative and ultimately detract from the model’s predictive ability. Thus, unlike other methods, the sequence processing and representation in `MHCrank` was designed to favor the amino acids near an antigen’s termini – it unifies the sequences of various lengths to length 10 (MHC Class I molecules favor peptides with 8-10 amino acids). Figure 1a presents the antigen representation process.

Specifically, if the peptide sequence is shorter than 10 amino acids, additional pseudo amino acids, represented by an ambiguous ’X’, are added at the center positions of the peptide (Figure 1a (A)). When the peptide’s original length is odd, one amino acid is padded offset towards theN-terminal side. This was motivated by past studies demonstrating that the C-terminal amino acids of a candidate antigen are more influential than the N-terminal amino acids with respect to whether or not a peptide will be processed for presentation.^2, 5^ If the peptide sequence is longer than 10 amino acids (Figure 1a (B)), a number of amino acids will be trimmed from the center of the peptide sequence, with one amino acid offset from the center towards the N-terminal if necessary.

### Amino acid representation

Once the peptide sequences are processed into uniform length, their remaining amino acids will be represented in various ways to capture the sequence contents. `MHCrank` has three distinct amino acid representation methods, denoted as `BLOSUM`, `embedding` and `em-BLO`, respectively. In the `BLOSUM` method, each amino acid is represented by its corresponding 21-dimension vector extracted from the BLOSUM62 substitution matrix.^17^ That is, the amino acid representation is hard coded as input to `MHCrank`. In the `embedding` method, each amino acid is first represented by an initial, random 21-dimension embedding vector; the embeddings will be learned^18^ in `MHCrank` so as to maximize their presentation power and facilitate optimal performance. 21 dimensions was selected to control for length given that this is also the same dimensions as the `BLOSUM` method. In the `em-BLO` method, an aggregate of both the non-learned vectors and learned embeddings (dimension 42) will be used. Note that the padded pseudo amino acid, X, is represented as a zero vector by `embedding`.

### MHCrank learning

Given the amino acid representations in a processed peptide, each peptide’s embedding matrix is further padded, following `MHCflurry-AP`,^8^ and forwarded to a 1-D convolutional layer with *n*_*k*_1__ kernels of size *k*, which aggregates different local information (*k*-mers) in the peptide. The output feature mapping is then forwarded to a number of components as described below that capture various signals from the training peptides and learn how each peptide’s flanking regions affect the probability that a peptide will undergo antigen processing.

#### Convolution over N-flank and C-flank

As was done in `MHCflurry`, the portion pertaining to the N-flank sequence is extracted from the feature mapping output of the first 1-D convolution layer. Mean pooling is conducted over the N-flank’s specific feature mapping to achieve the per-channel average for each amino acid in the N-flank. The results are then forwarded to a dense layer, which outputs a single value representing the flank’s favorability as a cleavage site. Identical operations are applied to the C-flank sequence.

#### Convolution over Peptide

Following `MHCflurry-AP`,^8^ convolutions are also applied to the peptide to learn the relationship between the cleavage favorability of its N-terminus/C-terminus and the cleavage favorability of its central amino acids. The intuition is that peptides with a higher cleavage favorability at their terminal position relative to their central amino acids are more likely to be processed for presentation than peptides in which this is not the case. To learn this relationship near the N-terminus, the portion pertaining to the peptide sequence is extracted from the feature mapping output of the first 1-D convolution layer and is subsequently forwarded to two stacked 1-D convolutional layers. The first layer has *n*_*k*_2__ channels; the second layer has 1 channel. Both of the stacked convolutional layers employ kernels of size 1. The output of the second layer contains a score for each amino acid in the peptide that represents the residue’s likelihood of being an N-terminal cleavage site. The scores corresponding to the N-terminal and C-terminal amino acids within the peptide (A in Figure 1b) are forwarded to the downstream learning. A max pool is applied over the non-N-terminal amino acids to identify the overall highest favorability for N-terminal cleavage to occur within the central amino acids. A max pool is also applied over the non-C-terminal amino acids to identify the overall highest favorability for C-terminal cleavage to occur within the central amino acids.

#### Convolution over C-terminal cleavage site

A novel component of `MHCrank` is the global-kernel based convolution over the C-terminal cleavage site. The C-terminal cleavage site is comprised of the terminal *r* amino acids of the peptide and the initial *r* amino acids of the C-flank, where *r* is the cleavage radius. The global kernel, referred to as a cleavage site-specific kernel and denoted as `CSSK`, is applied on amino acid representations of the cleavage site sequence to capture global signals useful for cleavage and processing. This was motivated by the fact that the C-terminal end of the peptide is more influential than the N-terminal with respect to cleavage and processing.^2, 5^ Because the C-terminus is the primary anchor point between the antigen and MHC molecule during binding, explicitly learning from the C-terminal cleavage site could enable additional, useful signals to predict a peptide’s favorability to be processed. The global kernel is used with the intuition that there are motifs located in this region, which are recognized by the peptidases and proteases that process peptides for presentation, that would be more easily learned and recognized by a global kernel.

#### Incorporating original peptide length

Compared to other methods, `MHCrank` has a novel, single node whose input is the peptide’s original length before padding or trimming. The downstream convolution’s use of the peptide’s original length in `MHCrank` is motivated by the fact that peptide length is a significant contributing factor to both processing and presentation.^1, 2^

#### Combing all information

Two fully-connected layers are applied in succession to the concatenated output of all the above components. Each layers possess *n*_*k*_2__ nodes. The output is then forwarded to an output layer that predicts the probability of a peptide undergoing antigen processing. Peptides more likely to undergo antigen processing receive higher probabilities.

### Ensemble methods and model selection

Ensembles^19^ have been an effective strategy in making more accurate predictions compared to that of a single model by reducing prediction variance. We developed the following six ensemble strategies to leverage and integrate multiple models, where the model performance on each fold was accessed using AUC score on each fold’s validation data, and overall model performance was accessed using the average AUC score across all four folds.

- `Fw-top1` (<u>f</u>old-<u>wi</u>se <u>top-1</u>): for each fold, we identified its best model and then combined these models. That is, the final ensemble consists of 4 total models – 1 from each of 4 folds. Please note that 4 models may correspond to different hyperparameters.
- `Fw-top2` (<u>f</u>old-<u>wi</u>se <u>top-2</u>): for each fold, we identified its top-2 best models and then combined these models. That is, the final ensemble consists of 8 total models – 2 from each of 4 folds. Please note that 8 models may correspond to different hyperparameters.
- `Ba-top1` (<u>b</u>est <u>a</u>verage <u>top-1</u>): we first identified the best performing hyperparameter set that had the best average AUC over the 4 folds. We then trained a model on each fold using the hyperparameter set. The final ensemble consists of 4 total models – 1 from each of 4 folds. Please note that these 4 models correspond to the same set of hyperparameters.
- `Ba-top2` (<u>b</u>est <u>a</u>verage <u>top-2</u>): we first identified the top-2 best performing hyperparameter sets that had the best average AUC over the 4 folds. We then trained a model on each fold using the hyperparameter set. The final ensemble consists of 8 total models – 2 from each of 4 folds. Please note that 4 models (1 from each of 4 folds) correspond to a single set of hyperparameters; the remaining 4 models (1 from each of 4 folds) correspond to another single set of hyperparameters. Furthermore, the set of hyperparameters belonging to the first 4 models is distinct from the set of hyperparameters belonging to the second 4 models.
- `C-top1` (<u>c</u>ombo <u>top-1</u>): we combined the `Fw-top1` and `Ba-top1` models. That is, the final ensemble consists of 8 total models, 4 from `Fw-top1` and 4 from `Ba-top1`.
- `C-top2` (<u>c</u>ombo <u>top-2</u>): we combined the `Fw-top2` and `Ba-top2` models. That is, the final ensemble consists of 16 total models, 8 from `Fw-top2` and 8 from `Ba-top2`.

The predicted scores from the ensemble models are calculated as the mean of the predicted scores from each of its component models. Table S1 outlines the specific hyperparameter sets of the models selected for each of the ensembles. The hyperparameters listed in Table S1 are only those that underwent tuning. The full set of hyperparameter options used to construct our `MHCrank` models can be found in Table S2. The training and validation AUC of the models selected for inclusion in each ensemble is detailed in Table S3.

### Model training

Three distinct model variations, each corresponding to one of the `embedding`, `BLOSUM`, `em-BLO` amino acid representation methods, combined with each hyperparameter set, were trained on each of the four training data folds. More information on the amino acid representations can be found in the ’Amino acid representation’ section. See ’Training data’ section for details regarding how the data was split into folds for training and validation. All hyperparameters utilized are listed in Table S2. Optimization was achieved using an Adam optimizer and a binary cross-entropy loss function. Models were trained for 500 epochs with an early stopping patience of 30 epochs.

### Evaluation metrics

Model were evaluated for ensemble selection by the AUC they achieved on validation folds. The fold and validation AUC for any model included in a `MHCrank` ensemble is presented in Table S3. `MHCrank` ensembles were compared to the `MHCflurry-AP` and `netMHCpan-EL` baselines via AUC, precision@*k*, and NDCG@*k* for *k* = {10, 25, 50, 100, 250, 500}.

- AUC: It is the area underneath a receiver operating characteristic (ROC) curve designed for binary classification problems. ROC a probability curve depicting true positive rates vs false positive rates over various prediction thresholds. Thus, AUC measures how well a model is capable of distinguishing between two classes (e.g., hits and decoys). Higher AUC values indicate better distinguishing capacity.
- Precision@*k*: It measures the proportion of top-*k* ranked peptides that are also hits. It is calculated as follows,

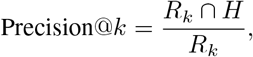

where *R_k_* is the set of top-*k* ranked peptides, and *H* is the set of hits. Higher precision@*k* scores indicate higher probabilities of correctly detecting hits within the top-*k* peptides. Note that precision@*k* is equivalent to the ‘PPV’ metric reported in O’Donnell *et al*.^8^
- NDCG@*k*: It is the normalized discounted cumulative gain (DCG) for top-*k* ranking. DCG@*k* is calculated as follows:

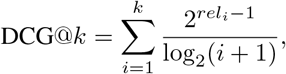

where *rel_i_* is the relevance of an peptide at position *i* indicating whether the recommended peptide is a hit (1) or a decoy (0); the numerator of the DCG@*k* equation awards relevant peptides and punishes decoys; the denominator gives more weight to higher ranked recommendations. NDCG@*k* is the normalized DCG@*k*. Higher NDCG@*k* scores indicates better performance.

### Statistical analysis

We obtained 1,000 sets of 100,000 peptides via bootstrap resampling of our testing data. Processing likelihood scores were obtained for each of these peptides from both our ensembles and the baseline methods. 95% confidence intervals (CIs) of our evaluation metrics were calculated to compare our ensemble models against both `MHCflurry-AP` and `netMHCpan-EL`. Non-overlapping CIs indicate a significant difference in performance at a threshold of at least *p*=0.05. Overlapping CIs indicate there was not a significant difference in performance at a significance level of *p*=0.05.

### Site-specific amino acid enrichment

A set of 50,000 randomly selected hits were obtained from the training peptides. For each model, the set of the top 100-recommended peptides were selected. The peptides in each set were processed to a length of 9 amino acid residues via the method outlined in Figure 1a and the ’Peptide preprocessing’ section. We produced a site-specific amino acid enrichment visualization from each set of peptides using the R package: `ggseqlogo`.^20^

### Cosine Similarity

Cosine similarity uses inner product space to measure the similarity between two vectors. The cosine similarity was calculated for the embedding vectors of each amino acid-pair and is given by the following:

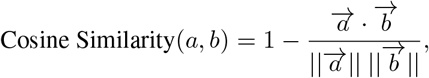

where 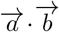 is the dot product of two vectors. Higher values indicate higher similarities.

## Experimental results

### Performance Evaluation

Tables 1, 2, and 3 present the performance of all six `MHCrank` ensembles, `MHCflurry-AP`, and `netMHCpan-EL`, in terms of mean AUC, precision@*k*, and NDCG@*k*, respectively. The corresponding 95% CIs used to measure significance (see Evaluation metrics section) are presented in Table S4. All (100%) of the models selected for inclusion in the ensemble utilized the `embedding` amino acid representation method (Table S1).

**Table 1:** Performance comparison of mean AUC

| MHCflurry-AP | netMHCpan-EL | MHCrank ensembles |  |  |  |  |  |
| --- | --- | --- | --- | --- | --- | --- | --- |
|  |  | Fw-top1 | Fw-top2 | Ba-top1 | Ba-top2 | C-top1 | C-top2 |
| 0.9106 | 0.9050 | 0.9073 | **\*0.9120** | 0.9102 | **\*0.9147** | **\*0.9121** | **\*0.9153** |

**Table 2:** Performance comparison of mean precision@*k*

| $k$ | MHCflurry-AP | netMHCpan-EL | MHCrank | | | | | |
| --- | --- | --- | --- | --- | --- | --- | --- | --- |
|  |  |  | Fw-top1 | Fw-top2 | Ba-top1 | Ba-top2 | C-top1 | C-top2 |
| 10 | 0.6717 | 0.7065 | **\*0.7260** | **\*0.7251** | 0.6492 | 0.6980 | **0.7089** | **0.7116** |
| 25 | 0.5901 | 0.6434 | **0.6544** | **0.6517** | 0.6314 | 0.6246 | **0.6496** | **0.6452** |
| 50 | 0.5482 | 0.5971 | **0.5973** | 0.5971 | 0.5891 | 0.5922 | **\*0.6063** | **0.6035** |
| 100 | 0.4827 | 0.5495 | 0.5284 | 0.5360 | 0.5182 | 0.5304 | 0.5491 | 0.5434 |
| 250 | 0.3994 | 0.4344 | 0.4202 | **\*0.4410** | 0.4198 | **0.4359** | **\*0.4412** | **\*0.4482** |
| 500 | 0.3215 | 0.3288 | **\*0.3413** | **\*0.3544** | **\*0.3409** | **\*0.3552** | **\*0.3550** | **\*0.3604** |

**Table 3:** Performance comparison of mean NDCG@*k*

| $k$ | MHCflurry-AP | netMHCpan-EL | MHCrank ensembles | | | | | |
| --- | --- | --- | --- | --- | --- | --- | --- | --- |
|  |  |  | Fw-top1 | Fw-top2 | Ba-top1 | Ba-top2 | C-top1 | C-top2 |
| 10 | 0.6884 | 0.7253 | **\*0.7451** | **\*0.7544** | 0.6705 | 0.7166 | **0.7265** | **0.7357** |
| 25 | 0.6230 | 0.6712 | **\*0.6846** | **\*0.6877** | 0.6486 | 0.6547 | **0.6761** | **0.6770** |
| 50 | 0.5802 | 0.6272 | **0.6317** | **0.6344** | 0.6113 | 0.6198 | **0.6346** | **0.6348** |
| 100 | 0.5179 | 0.5795 | 0.5658 | 0.5735 | 0.5486 | 0.5621 | **0.5805** | 0.5770 |
| 250 | 0.4328 | 0.4712 | 0.4202 | **\*0.4780** | 0.4538 | 0.4701 | **\*0.4772** | **\*0.4833** |
| 500 | 0.3545 | 0.3685 | **\*0.3779** | **\*0.3909** | **\*0.3743** | **\*0.3891** | **\*0.3909** | **\*0.3961** |

Table 1 shows that for all the methods, their mean AUC values were higher than 0.9. This suggests all methods have learned to distinguish between hits and decoys Among the evaluated methods, `C-top2` achieves the best performance (0.9153). All the `MHCrank` ensembles outperform `netMHCpan-EL`, and four of the six `MHCrank` ensembles – `Fw-top2`, `Ba-top2`, `C-top1`, and `C-top2` – outperform `MHCflurry-AP` with statistical significance. This demonstrates the strong power of `MHCrank` ensembles in learning from the training data to score hits above decoys. However, note that the high perceived accuracy as demonstrated by AUC may be optimistically inflated as a result of the class imbalance between hits and decoys in the test data. The AUC scores may indeed be inflated given the general reduction in relative performance for all methods when considering precision (Table 2) and NDCG (Table 3).

Table 2 illustrates that all `MHCrank` ensembles consistently outperform `MHCflurry-AP` across all *k* values with statistical significance. Furthermore, four of the six `MHCrank` ensembles (`Fw-top1`, `Fw-top2`, `C-top1` and `C-top2`) outperform `netMHCpan-EL` for small values of *k* (10, 25). For *k*=500, all `MHCrank` ensembles outperform `netMHCpan-EL` with a significance of at least *p* = 0.05. Among the six `MHCrank` ensemble methods, `Ba-top1` and `Ba-top2` are generally the worst performing ensembles in terms of precision. On average, neither ensemble improves upon the performance of `netMHCpan-EL`. Furthermore, `Ba-top1` is the only ensemble to achieve a significantly worse precision than `MHCflurry-AP` for any value of *k* (10). These two ensemble methods used the overall best hyperparameters across all the 4 folds. Thus, the models trained on each fold using these hyperparameter sets were not necessarily optimized for that fold. Consequently, the combination of these sub-optimal models did not produce the best performance. On the contrary, `Fw-top2` and `C-top1` were the two ensemble methods that achieved the overall best precision. Both methods incorporated at least the best model (`C-top1`) or two best models (`Fw-top2`) for each of the 4 folds, allowing the ensemble to integrate the most predictive power possible from the data.

Table 3 displays very similar trends to those in Table 2. That is, all the `MHCrank` methods outperform `MHCflurry-AP`, and four of the six ensemble methods (`Fw-top1`, `Fw-top2`, `C-top1` and `C-top2`) outperform `netMHCpan-EL`. Again, `Ba-top1` and `Ba-top2` are the worst performing ensemble methods. Unlike Table 2, `Fw-top1` and `Fw-top2` are the two best performing ensembles, with each achieving significant improvement over `netMHCpan-EL` for 3 and 4 values of *k*, respectively. This indicates that `Fw-top2` is able to rank more hits at higher positions in the ranking. `C-top1` and `C-top2` combine `Fw-top1` and `Ba-top1`, and `Fw-top2` and `Ba-top2`, respectively. This grants `C-top1` and `C-top2` both pros and cons of both types of ensembles. Thus it is intuitive that they, in consequence, achieved mid-level performance.

Table 4 summarizes the performance improvement of `MHCrank` as a percentage of its best performing ensemble: `Fw-top2`, over `MHCflurry-AP` and `netMHCpan-EL` on mean AUC, precision@*k* and NDCG@*k*. For both precision and NDCG, the percent improvement garnered by `Fw-top2` over both `netMHCpan-EL` and `MHCflurry-AP` exhibits generally increasing performance with increasing values of *k* (e.g., *k*=10 vs *k*=500). All percent improvements over `MHCflurry-AP` are significant for both metrics. This again indicates `Fw-top2` is able to more effectively rank peptides likely to be processed for presentation among the very top of ranking lists when compared to `netMHCpan` and `MHCflurry`. One surprising result is the dip in both precision and NDCG of `Fw-top2` relative to `netMHCpan-EL` for *k*=50, 100, followed by rapid improvement between *k*=250, 500. One plausible explanation is `netMHCpan-EL` learned to highly rank peptides with a specific motif that is highly enriched within hits (Figure 2). After exhausting all peptides containing the learned motif, the accuracy of subsequently ranked peptides would likely deteriorate.

**Table 4:**
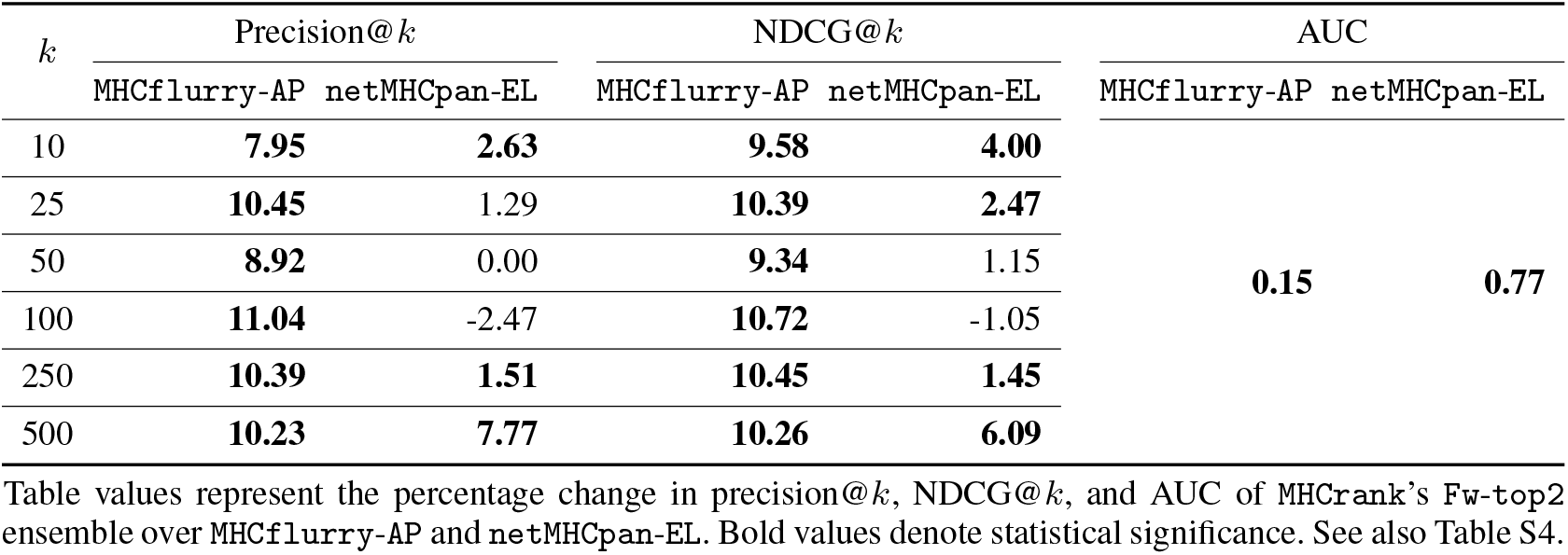
Performance comparison (% change)

**Figure 2:**
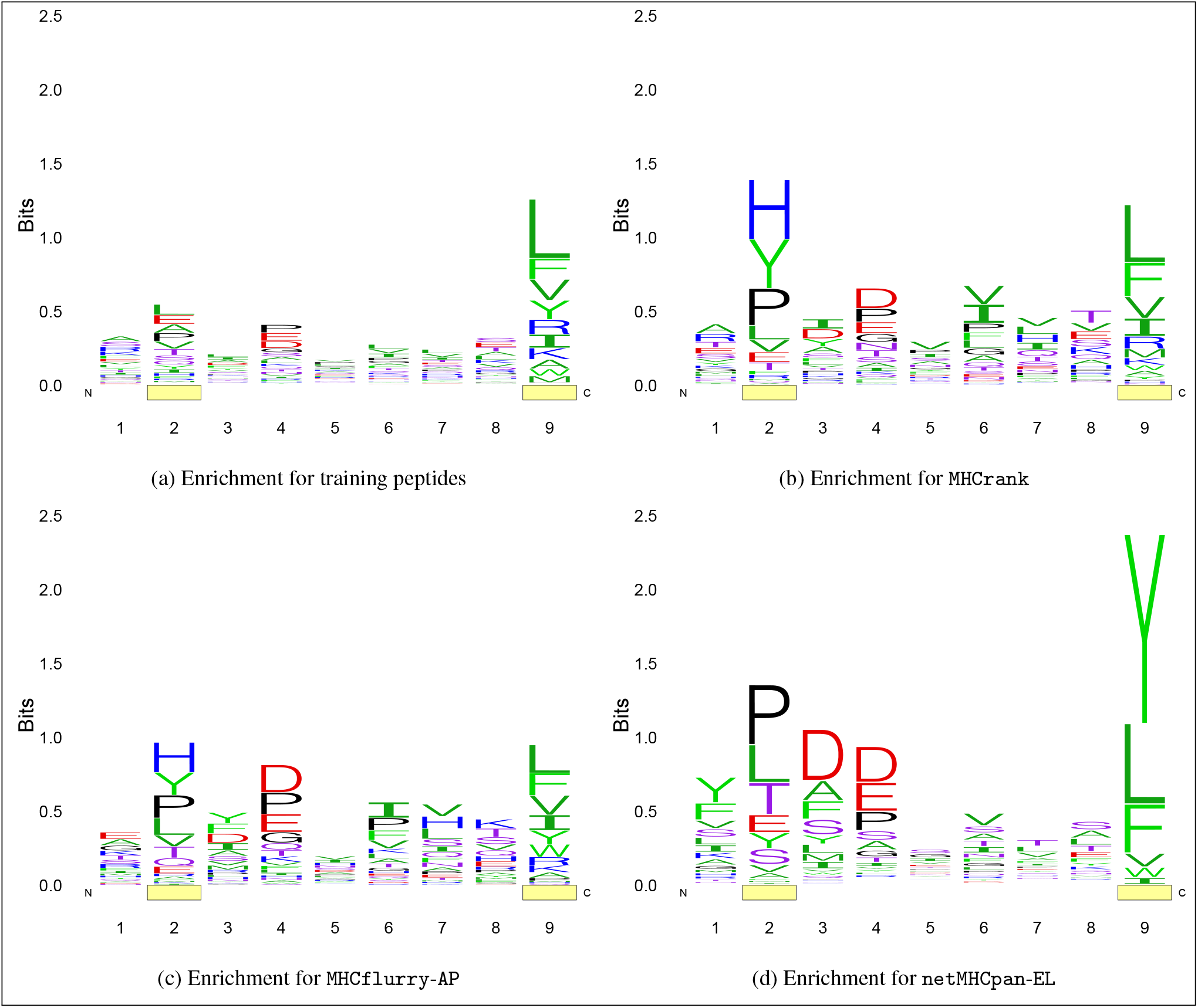
Position specific amino acid enrichment. Enrichment of amino acids in (**2a**) 50,000 randomly sampled hits from training data set and in the top 100 peptides from the testing data ranked by (**2b**) `MHCrank`’s `Fw-top2` ensemble and both the (**2c**) `MHCflurry-AP` and (**2d**) `netMHCpan-EL` baseline methods. Yellow boxes covering positions 2 and 9 in each figure highlights the enrichment of the peptides at their typical anchor positions.

### Enrichment of top-ranked peptides

Figure 2 shows the position-specific enrichment in various sets of peptides. The 2^nd^ and 9^th^ positions are underscored in each figure with yellow boxes. These positions correspond to the canonical anchor residues for binding to the MHC Class I molecule. Figure 2a illustrates the position-specific enrichment for 50,000 randomly selected hits from the training data. In this figure, position 9 exhibits a high level of enrichment. The amino acids that are enriched at this position are either hydrophobic (L, V, I, etc.) or aromatic (F, Y, W) and are all enriched to comparable levels. This conforms to the biological relevance affirmed by previous studies^3^ that hydrophobic residues tend to be favored in the C-terminal position. Position 2 also has slightly elevated enrichment when compared to the low levels of enrichment that exist for positions 1 and 3 – 8. Taken together, this demonstrates that the peptides selected for training can represent a large range of different peptides.

Figure 2b depicts the position-specific enrichment for the set of the top 100-ranked peptides by the best `MHCrank` ensemble `Fw-top2`. The top 100 peptides were selected for each model as *k*=100 is the threshold where `MHCrank`’s performance rapidly improves compared to `netMHCpan-EL`. By examining the general composition of peptides ranked highly by `MHCrank` and the `netMHCpan-EL` and `MHCflurry-AP` baselines, we believe we may be able to elucidate the reason for the drastic shift in performance at *k*=100 that we observed in Table 4. The position-specific enrichment of `Fw-top2`’s recommended peptides shows an enhancement in the enrichment at the 2^nd^ and 9^th^ positions relative to Figure 2a. Their enhancement in the `Fw-top2` ensemble’s top-100 ranked peptides indicates that our method is capable of discerning both anchor positions within the peptides. Moreover, similar to the enrichment of the training peptides, the patent enrichment at the 9^th^ position features uniform enrichment of mostly hydrophobic amino acids. This suggests that `Fw-top2` learned to identify the features and physiochemical properties of the residues versus specific sequences. Note there was also a slight increase in the enrichment of the central amino acids (positions 3-9) compared to Figure 2a, indicating that `MHCrank` may have learned some motif(s) that convey processing favorability within the central amino acids.

Figure 2c depicts the position-specific enrichment for the set of top-100 ranked peptides by `MHCflurry-AP`. Like `MHCrank`, `MHCflurry-AP` exhibited enhanced enrichment of the 2^nd^ position. However, this is overshadowed by similar enrichment levels of its central amino acids (positions 3-9). Additionally, Figure 2c displays a reduction in enrichment for the vital C-terminal position. Thus, It appears there was not any position for which `MHCflurry-AP` was able to learn meaningful features nor trends. This lack of fit might explain why `MHCflurry-AP`’s performance was worse than `MHCrank`’s performance.

Figure 2d depicts the position-specific enrichment for the set of the top-100 ranked peptides by `netMHCpan-EL`. The enrichment of the 9^th^ position is higher than any positionspecific enrichment from both `MHCrank` and `MHCflurry-AP`. Unlike the enrichment at this position present in `MHCrank`’s top peptides, only three amino acids (Y, L, F) are enriched for `netMHCpan-EL`. This is an important distinction because rather than learning properties of the amino acids that occupy this position in hits, it is likely that `netMHCpan-EL` learned to prefer peptides that ended in one of the three enriched amino acids. In fact, 91% of `netMHCpan-EL`’s top-100 peptides feature either Y, L, or F in the C-terminal position, suggesting that `netMHCpan-EL`’s performance may decline when testing it with peptides that do not match this pattern. This adds credence to our explanation underlying the dip in both precision and NDCG of `Fw-top2` relative to `netMHCpan-EL` for *k*=50, 100, followed by rapid improvement between *k*=250, 500 (Table 4). Also note `netMHCpan-EL`’s enhanced enrichment at positions 2, 3, and 4. The enrichment of positions 2-4 suggests `netMHCpan-EL` was able to learn that there is an important feature near those positions, but not which position was most informative.

### Similarities of learned amino acid embeddings

We extracted the 21-dimension embedding vector for each amino acid from a representative `MHCrank` model. Figure 3 illustrates the cosine similarities among the embeddings learned by `MHCrank` for each amino acid. Amino acids in Figure 3 have been grouped according to their types: hydrophobic, aromatic, basic, acidic, polar, and other. We observed that, in general, the learned embeddings of amino acids within the same groups (i.e., of the same types) are more similar than those of amino acids from different groups (i.e., of different types). This indicates that `MHCrank` was capable of learning meaningful information from amino acids that may correlate with their physiochemical properties, and thus facilitate better predictions. This is further demonstrated by the similarities of learned embeddings of amino acids between certain groups. Figure 3 shows that the embeddings of aromatic and hydrophobic amino acids are more similar to each other than the other amino acid types. Likewise, the embeddings of basic, acidic, and polar amino acids are more similar to each other than they are to other amino acid types. This distinction suggests that `MHCrank` is capable of learning information that corresponds to an amino acid’s hydrophilicity, an important physiochemical property involved in identifying peptides likely to be processed.

**Figure 3:**
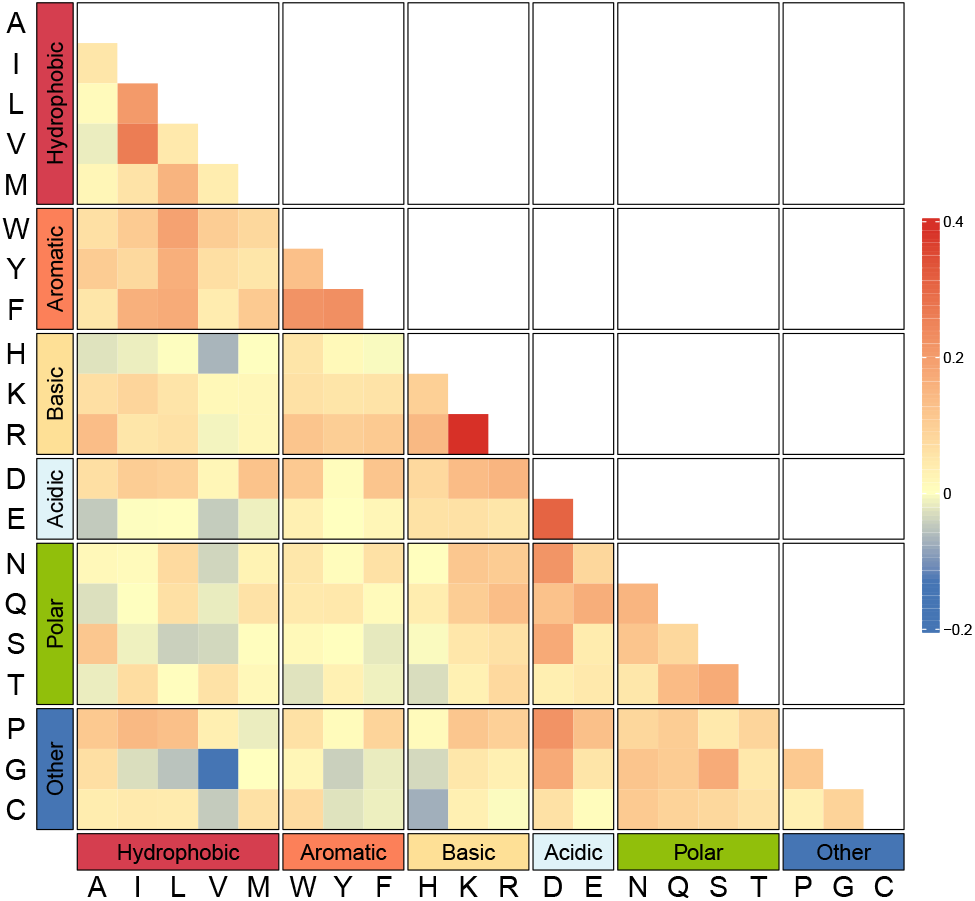
Cosine similarity matrix of learned amino acid embeddings

## Discussion

We observed that the all but two of the models included in the `MHCrank` ensembles processed peptides to a length of 9 or 10 amino acids, and all models utilized the `embedding` amino acid representation. These findings demonstrate that peptide representations are improved through the enhanced biological relevance of our pre-processing method and the inclusion of learned embeddings. For pre-processing, we believe that our leveraging the knowledge that central amino acids of longer peptides often do not interact with the MHC Class I molecule enabled processed peptides to retain only the most relevant information. The dearth of enrichment in the central amino acids both in the training peptides (Figure 2a) and in the top-100 ranked peptides from `MHCrank` (Figure 2b), paired with our improved performance over `MHCflurry-AP` (Table 4), further strengthens the claim that central amino acids are not necessarily relevant features and, their removal can improve model performance.

With respect to the unique embedding learned for each amino acid, the enrichment we observed in Figure 2 also highlights their capability to identify features and information that may not have been imparted by the `BLOSUM` matrix. As observed in Figure 2b, all but one of the hydrophobic amino acids are enriched to comparable levels at the C-terminal position in `Fw-top2`’s top predicted peptides. Not only does this coincide with biological observations, but as we observed in Figure 3, it also suggests that `MHCrank` has learned to identify, *and favor*, certain physiochemical properties of amino acids, such as hydrophobicity, despite no *a priori* knowledge.^1^ Furthermore, the relative enrichment of `MHCrank`’s 9^th^ position versus its 7^th^ and 8^th^ positions (Figure 2b), suggests that the implementation of the `CSSK` aids in the identification of commonalities among protease cleavage site motifs. This is especially apparent when considering the enrichment of the same positions from peptides ranked by `MHCflurry-AP`.

Thus, even for different numbers of top-*k* ranked peptides based on predictions, where the performance of `MHCrank` and `netMHCpan` were not significantly different, we believe `MHCrank` might still be considered superior as it achieves better performance using more unique peptides. `MHCrank` also achieved superior performance compared to `MHCflurry-AP` for all evaluated metrics. In addition, the amino acid learned embeddings and enrichment was highly correlated with biological observations. Altogether, the proposed `MHCrank` demonstrates strong performance compared to existing methods, and could have vast applicability to aid drug and vaccine development.

### Limitations and future directions

Despite `MHCrank`’s strong performance, there are multiple improvements that might further strengthen its relevance and applicability for drug and vaccine development. First, the implementation of a combinatorial approach that incorporates binding affinity as well – similar to O’Donnell *et al*’s presentation score predictor^8^ – might improve upon the current presentation predictions achieved for the MHC class I molecule.

Secondly, given that the purpose of predicting MHC Class I presentation is to aid in the development of drugs and vaccines that stimulate the adaptive immune response through the activation of T-cells, future endeavors may benefit from creating models that predict both the magnitude and type of response a specific antigen will elicit. For MHC Class I presented antigens in particular, this might be accomplished by training models to identify complementary sequences between the presented peptide’s central amino acids and the T-cell receptor’s (TCR) active site to which it binds.

A final limitation of not only of `MHCrank`, but also of other MHC binding prediction models, is their inability to effectively utilize protein structure for predictions. While BLOSUM62^17^ and other popular embedding schemes aim to encapsulate sequence homology and dissimilarities or similarities between amino acids, this is a sub-optimal approach. The sequence of a peptide is not, in and of itself, deterministic of an antigen’s ability to be processed, presented, and recognized. Rather, it is a product of physiochemical properties of individual amino acids interacting with one another.^3^ The utilization of structural data requires a structure to first be resolved in a lab setting, rendering the approach infeasible. However, as structural prediction algorithms improve and become increasingly biologically relevant, abandoning sequences for structures will likely improve model performance. Specifically, the use of geometric deep learning models seem poised to yield the highest probability of success.^21^ We will investigate along these lines in our future research.

## Reproducibility

### Data and code availability

`MHCrank` source code, training and testing data as well as the trained neural networks used to create `MHCrank` ensembles are freely available here: https://github.com/ninglab/mhcrank.

### Computing resources

Data processing and model training, validation, and testing were all executed on Pitzer clusters of the Ohio Supercomputer Center.^22^ We implemented models using Python-3.6.6 and TensorFlow-2.2.1. We trained models with 1 Intel Xeon 8268s Cascade Lakes CPU node and 1 NVIDIA Volta V100 GPU totaling 32 GB of memory.

### Hyperparameters

A total of 729 models were trained across each of the 4 folds and evaluated. Each model was produced using a unique combination of hyperparameters. The specific hyperparameter options evaluated is presented below in Table S2. Note that while the table presents only 243 unique combinations, these would be applied to each of the 3 amino acid representation methods: `embedding`, `BLOSUM`, and `em-BLO`.

## Acknowledgments

This project was made possible, in part, by support from the National Institute of General Medical Sciences (2R01GM118470-05) and AWS Machine Learning Research Award. Any opinions, findings, and conclusions or recommendations expressed in this material are those of the authors, and do not necessarily reflect the views of the funding agencies.

## Author contributions

X.N. conceived the research, obtained funding for the research, and supervised P.L.; P.L. and X.N. designed the research; P.L. and X.N. conducted the research, including data curation, formal analysis, methodology design and implementation, visualization; P.L. drafted the original manuscript; P.L. and X.N. conducted the manuscript editing P.L. and X.N. reviewed the manuscript.

## Supplementary Materials

**Table S1:** Hyperparameters for each of the best performing `MHCrank` models selected for inclusion in ensembles

| Ensemble method | Fold | A.A. | peptide length | $r$ | $k$ | $n_{k_1}$ | $n_{k_2}$ |
| --- | --- | --- | --- | --- | --- | --- | --- |
| Fw-top1 | 0 | embedding | 10 | 2 | 9 | 512 | 64 |
|  | 1 | embedding | 10 | 2 | 11 | 512 | 32 |
|  | 2 | embedding | 10 | 2 | 11 | 512 | 64 |
|  | 3 | embedding | 15 | 2 | 11 | 512 | 64 |
| Fw-top2 | 0 | embedding | 9 | 2 | 11 | 512 | 64 |
|  | 1 | embedding | 10 | 1 | 11 | 512 | 32 |
|  | 2 | embedding | 10 | 2 | 11 | 512 | 32 |
|  | 3 | embedding | 15 | 1 | 11 | 512 | 64 |
| Ba-top1 | - | embedding | 10 | 2 | 9 | 512 | 64 |
| Ba-top2 | - | embedding | 10 | 2 | 11 | 512 | 64 |

A.A. refers to the amino acid representation method used by the model. Additionally, peptide length is the number of amino acids to be included in the processed peptide; $r$ is the cleavage radius; $k$ denotes kernel size; $n_{k_1}$ is the number of filters employed in convolutional layers; and $n_{k_2}$ is the dimension of the final two dense layers. Fw-top2 includes those listed as well as those in Fw-top1. Likewise, Ba-top2 includes the models possessing the hyperparameters from Ba-top1. Because Ba-top1 and Ba-top2 use the models trained on each of the folds for the listed, no folds are listed.

**Table S2:** Hyperparameter options employed by `MHCrank`

| Hyperparameter | Values |
| --- | --- |
| Peptide length | 9, 10, 15 |
| Flank length | 5 |
| $r$ | 1, 2, 3 |
| $n_{k_1}$ | 128, 256, 512 |
| $k$ | 7, 9, 11 |
| Convolution activation function | ReLU |
| $\ell_1$ $\ell_2$ regularization | [0.0, 0.0] |
| $n_{k_2}$ | 16, 32, 64 |
| Dropout rate | 0.5 |
| Loss function | Binary cross-entropy |
| Optimizer | Adam |
| Learning rate | 0.001 |

### MHCrank training results

Table S3 presents the training and validation AUC for any models included in any of the `MHCrank` ensembles (see ’Ensemble methods and model selection’ section). We observe a decrease in AUC from training to validation for all models. This is expected as models will have learned information from the peptides in the training set, while the peptides in the validation set can be considered novel. Interestingly, we observed larger drops in AUC from the training to the validation set for the two best performing models of those trained on fold 3 compared to the top performing models from other folds. Furthermore, the best performing model of those trained on fold 3 exhibits a large standard deviation in both mean training and validation AUC compared to all other selected models. This is because the model with this set of hyperparameters achieved poor performance when trained on fold 0. This suggests that the hyperparameters used by this model are not conducive to learning the important features of fold 0’s data. Moreover, the only hyperparameter that is shared by the best performing models from fold 3 and not present in any of the other top models is that those from fold 3 use a input peptide length of 15. This indicates that perhaps a larger proportion of peptides in fold 3 have central amino acids that contribute to MHC Class I processing or binding than those of the other folds.

**Table S3:** Training and Validation AUC of models included in `MHCrank` ensembles

| Ensemble | Model | Fold 0 |  | Fold 1 |  | Fold 2 |  | Fold 3 |  | Mean (STD) |  |
| --- | --- | --- | --- | --- | --- | --- | --- | --- | --- | --- | --- |
|  |  | Train | Val | Train | Val | Train | Val | Train | Val | Train | Val |
| Fw-top1 | fold0<sub>1</sub> | **0.9371** | **\*0.8119** | 0.9298 | 0.8249 | 0.9428 | 0.8354 | 0.9467 | 0.7986 | 0.9391 (7.4e-3) | 0.8177 (1.6e-2) |
|  | fold1<sub>1</sub> | 0.9476 | 0.7858 | **0.9462** | **0.8304** | 0.9451 | 0.8362 | 0.9507 | 0.7937 | 0.9459 (4.4e-3) | 0.8115 (2.5e-2) |
|  | fold2<sub>1</sub> | 0.9469 | 0.8039 | 0.9492 | 0.8269 | **0.9515** | **\*0.8433** | 0.9518 | 0.7939 | 0.9499 (2.3e-3) | 0.8170 (2.2e-2) |
|  | fold3<sub>1</sub> | 0.5000 | 0.5000 | 0.9308 | 0.8141 | 0.9353 | 0.8287 | **0.9522** | **0.8012** | 0.8296 (2.2e-1) | 0.7360 (1.6e-1) |
| Fw-top2 | fold0<sub>2</sub> | **0.9471** | **0.8092** | 0.9434 | 0.8219 | 0.9382 | 0.8272 | 0.9530 | 0.7956 | 0.9454 (6.2e-3) | 0.8134 (1.4e-2) |
|  | fold1<sub>2</sub> | 0.9407 | 0.7998 | **0.9432** | **0.8302** | 0.9489 | 0.8314 | 0.9399 | 0.7975 | 0.9432 (4.1e-3) | 0.8147 (1.9e-2) |
|  | fold2<sub>2</sub> | 0.9476 | 0.7858 | 0.9462 | 0.8304 | **0.9451** | **0.8362** | 0.9507 | 0.7937 | 0.9459 (4.4e-3) | 0.8115 (2.5e-2) |
|  | fold3<sub>2</sub> | 0.9384 | 0.7980 | 0.9479 | 0.8255 | 0.9528 | 0.8346 | **0.9472** | **0.7988** | 0.9465 (6.0e-3) | 0.8142 (1.9e-2) |
| Ba-top1 |  | **0.9371** | **\*0.8119** | **0.9298** | **0.8249** | **0.9428** | **0.8354** | **0.9467** | **0.7986** | 0.9391 (7.4e-3) | 0.8177 (1.6e-2) |
| Ba-top2 |  | **0.9469** | **0.8039** | **0.9492** | **0.8269** | **0.9515** | **\*0.8433** | **0.9518** | **0.7939** | 0.9499 (2.3e-3) | 0.8170 (2.2e-2) |

Values represent the training (Train) and validation (Val) AUC calculated for the model created by training a distinct hyperparameter set on each fold. Mean (STD) is calculated by averaging the AUC for all 4 models obtained from training a single hyperparameter set on each fold. Bold values indicate which models were included in the ensembles. A "\*" next to a selected model indicates that the model was included in multiple ensembles. To that end, Ba-top1 is the same set of hyper-parameters as fold0<sub>1</sub> and Ba-top2 is the same set of hyper-parameters as fold2<sub>1</sub>.

**Table S4:** Mean precision and NDCG@*k* and AUC (± a 95% CI)

| $k$ | MHCflurry-AP | | netMHCpan-EL | |
| --- | --- | --- | --- | --- |
| | precision@$k$ | NDCG@$k$ | precision@$k$ | NDCG@$k$ |
| 10 | 0.6717±9.2e-3 | 0.6884±9.9e-3 | 0.7065±9.0e-3 | 0.7253±9.3e-3 |
| 25 | 0.5901±6.1e-3 | 0.6230±6.5e-3 | 0.6434±5.8e-3 | 0.6712±6.2e-3 |
| 50 | 0.5482±4.4e-3 | 0.5802±4.8e-3 | 0.5971±4.3e-3 | 0.6272±4.6e-3 |
| 100 | 0.4827±3.2e-3 | 0.5179±3.5e-3 | 0.5495±3.2e-3 | 0.5795±3.4e-3 |
| 250 | 0.3994±1.9e-3 | 0.4328±2.1e-3 | 0.4344±2.1e-3 | 0.4712±2.2e-3 |
| 500 | 0.3215±1.4e-3 | 0.3545±1.5e-3 | 0.3288±1.4e-3 | 0.3685±1.5e-3 |
| AUC | 0.9106±.7e-4 | | 0.905±3.8e-4 | |
| $k$ | MHCrank Fw-top1 | | MHCrank Fw-top2 | |
| | precision@$k$ | NDCG@$k$ | precision@$k$ | NDCG@$k$ |
| 10 | 0.7260±9.0e-3 | 0.7451±9.2e-3 | 0.7251±9.3e-3 | 0.7544±9.3e-3 |
| 25 | 0.6544±6.2e-3 | 0.6846±6.4e-3 | 0.6517±6.1e-3 | 0.6877±6.3e-3 |
| 50 | 0.5973±4.4e-3 | 0.6317±4.6e-3 | 0.5971±4.3e-3 | 0.6344±4.5e-3 |
| 100 | 0.5284±3.2e-3 | 0.5658±3.4e-3 | 0.5360±3.2e-3 | 0.5735±3.3e-3 |
| 250 | 0.4202±1.9e-3 | 0.4596±2.1e-3 | 0.4410±2.0e-3 | 0.4780±2.1e-3 |
| 500 | 0.3413±1.3e-3 | 0.3779±1.4e-3 | 0.3544±1.4e-3 | 0.3909±1.5e-3 |
| AUC | 0.9073±4.0e-4 | | 0.9120±4.0e-4 | |
| $k$ | MHCrank Ba-top1 | | MHCrank Ba-top2 | |
| | precision@$k$ | NDCG@$k$ | precision@$k$ | NDCG@$k$ |
| 10 | 0.6492±9.3e-3 | 0.6705±1.0e-2 | 0.6980±9.3e-3 | 0.7166±9.6e-3 |
| 25 | 0.6314±6.0e-3 | 0.6486±6.5e-3 | 0.6246±6.1e-3 | 0.6547±6.5e-3 |
| 50 | 0.5891±4.3e-3 | 0.6113±4.7e-3 | 0.5922±4.3e-3 | 0.6198±4.7e-3 |
| 100 | 0.5182±3.0e-3 | 0.5486±3.7e-3 | 0.5304±3.0e-3 | 0.5621±3.3e-3 |
| 250 | 0.4198±2.0e-3 | 0.4538±2.1e-3 | 0.4359±2.0e-3 | 0.4701±2.1e-3 |
| 500 | 0.3409±1.4e-3 | 0.3743±1.5e-3 | 0.3552±1.4e-3 | 0.3891±1.5e-3 |
| AUC | 0.9102±4.0e-4 | | 0.9147±3.8e-4 | |
| $k$ | MHCrank C-top1 | | MHCrank C-top2 | |
| | precision@$k$ | NDCG@$k$ | precision@$k$ | NDCG@$k$ |
| 10 | 0.7089±9.0e-3 | 0.7265±9.5e-3 | 0.7116±9.4e-3 | 0.7357±9.5e-3 |
| 25 | 0.6496±6.0e-3 | 0.6761±6.5e-3 | 0.6452±6.0e-3 | 0.6770±6.3e-3 |
| 50 | 0.6063±4.4e-3 | 0.6346±4.7e-3 | 0.6035±4.4e-3 | 0.6348±4.6e-3 |
| 100 | 0.5491±3.1e-3 | 0.5805±3.4e-3 | 0.5434±3.1e-3 | 0.5770±3.3e-3 |
| 250 | 0.4412±2.0e-3 | 0.4772±2.1e-3 | 0.4482±2.0e-3 | 0.4833±2.1e-3 |
| 500 | 0.3550±1.4e-3 | 0.3909±1.5e-3 | 0.3604±1.4e-3 | 0.3961±1.5e-3 |
| AUC | 0.9121±3.9e-4 | | 0.9153±3.8e-4 | |

